# CXCL12/CXCR4 Governs Lymphatic Valve Formation Through Flow Dependent AKT/FOXO1/FOXC2 Activation

**DOI:** 10.64898/2026.07.21.739861

**Authors:** Long Nguyen Hoang Do, Jingjing Pang, Esteban Delgado, Liam Flynn, Hongxia Liu, Xiaolei Liu

## Abstract

**Background:** Lymphatic valves are specialized structures within lymphatic vessels that ensure unidirectional lymph transport. Defective lymphatic valve formation is associated with lymphedema, a chronic disease characterized by impaired lymph drainage and the accumulation of protein-rich interstitial fluid. Lymphatic valves form during late embryonic stages in mice, with oscillatory shear stress implicated in regulating the molecular mechanisms which control lymphatic valve formation. However, the molecular mechanisms governing lymphatic valve formation remain incompletely understood. Although CXCR4 is regulated by shear stress in blood vessels, whether CXCL12/CXCR4 signaling regulates lymphatic valve formation and the underlying molecular mechanisms remain unknown.

**Methods:** To investigate the roles of CXCR4 in the regulation of lymphatic valve development, we utilized lymphatic endothelial cell (LEC)–specific *Cxcr4* knockout (*Flt4CreER^T2^, Cxcr4^f/f^*) mice. To determine the source of CXCL12, major ligand for CXCR4 in the mesentery, we used global *Cxcl12*−/−, and *Cxcl12-DsRed* knock-in/knockout (KIKO) reporter mice, as well as conditional *Cxcl12* knockout mouse lines. To determine the molecular mechanisms by which CXCL12/CXCR4 regulates lymphatic valve development, primary human dermal LECs were exposed to oscillatory shear stress (OSS) to mimic valve-associated flow, followed by analysis of downstream signaling pathways and valve related gene expression.

**Results:** LEC-specific loss of CXCR4 displayed impaired lymphatic valve development. *Flt4CreER^T2^, Cxcr4^f/f^* mice showed a significant reduction in valve numbers in embryonic mesenteric lymphatic vessels. Similarly, reduced valve numbers were observed in *Cxcl12−/−* embryos, indicating CXCL12/CXCR4 is required for embryonic mesentery collecting lymphatic valve formation. *Cxcl12-DsRed* KIKO mice revealed that blood vessels, opposed to nerves, were the major source of CXCL12 in the embryonic mesentery. In align with this finding, EC-specific *Cxcl12* deletion recapitulated defective valve phenotypes observed in *Cxcl12*−/− embryos. Mechanistically, *CXCR4* knockdown in primary human dermal LECs attenuated OSS induced phosphorylation of AKT and FOXO1, leading to increased nuclear localization of FOXO1 and reduced expression of FOXC2, an essential transcription factor governing lymphatic valve development. Consistent with these findings, lymphatic valves of LEC-*Cxcr4* deficient mice exhibited increased FOXO1 nuclear localization. Importantly, pharmacological activation of AKT reduced FOXO1 nuclear accumulation and restored lymphatic valve numbers in LEC-*Cxcr4* deficient mesenteric lymphatic vessels.

**Conclusions:** Our findings reveal CXCL12/CXCR4 signaling acts as a critical regulator of lymphatic valve development. CXCL12/CXCR4 signaling integrates into the flow dependent AKT/FOXO1/FOXC2 signaling axis to coordinate lymphatic valve development and morphogenesis. Taken together, our study uncovers a previously unknown role of CXCL12/CXCR4 signaling pathway in the regulation of lymphatic valve development. Targeting CXCL12/CXCR4/AKT/FOXO1 axis may represent a promising therapeutic strategy for improving lymphatic valve development to improve lymphatic function for the treatment of lymphedema.

## Introduction

The lymphatic vasculature plays an essential role in maintaining tissue fluid homeostasis, mediating immune cell uptake and transport, and facilitating dietary lipid absorption ^1^. Efficient lymph transport depends on the presence of lymphatic valves, specialized intraluminal structures within collecting lymphatic vessels that prevent retrograde flow ensuring unidirectional lymph movement ^2^. Defects in lymphatic valve development or function disrupt lymphatic circulation and contribute to lymphatic disorders such as lymphedema ^3^. Lymphatic valve formation is initiated during embryogenesis at sites of disturbed or oscillatory lymph flow, where LECs sense biomechanical forces to undergo coordinated transcriptional and morphological changes generating valve leaflets ^4,5^. This process is governed by a conserved transcriptional network including prospero homeobox 1 (PROX1), forkhead box C2 (FOXC2), and GATA-binding protein 2 (GATA2), which together regulate the expression of valve-associated genes and lymphatic valve development ^6–8^. In addition to transcriptional regulation, mechanotransduction pathways play a central role in valve morphogenesis, as LECs translate oscillatory shear stress into signaling cascades that coordinate endothelial alignment, clustering, and leaflet formation ^4,9^. Despite these advances in our knowledge, the molecular mechanisms that integrate biochemical signaling with flow-dependent mechanotransduction to control lymphatic valve development remain incompletely understood.

Emerging evidence indicates that chemokine signaling pathways contribute to lymphatic vascular development and endothelial cell behavior. In particular, the CXCL12/CXCR4 axis regulates lymphatic endothelial migration, survival, and vascular patterning ^10–13^. CXCL12/CXCR4 signaling activates downstream pathways including PI3K/AKT ^13,14^. Knockdown of CXCR4 in LECs inhibits VEGFC/ VEGFR3 induced PI3K/AKT signaling activation and disrupts transcriptional programs that govern endothelial remodeling ^15^. A key downstream target of AKT in endothelial cells is the transcription factor FOXO1, a repressor of valve-forming genes whose activity is inhibited by AKT-mediated phosphorylation and subsequent nuclear exclusion ^16–17^. However, whether CXCL12/CXCR4 signaling directly regulates lymphatic valve formation, and how it connects with AKT/FOXO1 signaling during this process, remain unclear.

Here, we demonstrate that CXCL12/CXCR4 signaling is required for lymphatic valve formation and collecting vessel maturation, acting through a signaling network that regulates AKT/FOXO1 activity in LECs. And we found that mesenteric blood vessels are a major source of CXCL12 and show that endothelial cell derived CXCL12 signals through CXCR4 on LECs to drive lymphatic valve formation. These findings identify a previously unrecognized mechanism linking chemokine signaling and flow dependent cues to control lymphatic valve development and provide new insight into the molecular regulation of lymphatic vascular morphogenesis.

## Materials and Methods

### Animals and Treatments

Mouse strains used in this study included *Flt4CreER^T2^* ^18^, *Cxcr4^f/f^* (Jackson Laboratory, stock no. 008767), *Cxcl12-DsRed* knock-in/knockout (KIKO) reporter mice (Jackson Laboratory, stock no. 022458), and *Cdh5CreER^T2^* mice have been previously described ^19^. All mice were maintained on a mixed C57BL/6J and NMRI genetic background. For embryonic deletion of *Cxcr4*, pregnant dams carrying *Flt4-CreER^T2^*; *Cxcr4^f/f^* (embryos received two consecutive intraperitoneal injections of tamoxifen (Sigma, T5648) at a dose of 5 mg/40 g body weight at the embryonic (E) 13.5 and half dose at E14.5, as described in the results section. For AKT activation experiments, the AKT activator SC-79 (Sigma, 123871) was administered 40 mg/kg to pregnant dams carrying embryos at E15.5. All animal experiments and husbandry procedures were performed in accordance with protocols approved by the Temple University Institutional Animal Care and Use Committee.

### Cell culture and Treatments

Human dermal LECs were purchased from Promocell (C-22121) and cultured with endothelial basal medium complemented with supplement mix (Promocell, C-39221) and 5% penicillin/streptomycin, as reported previously ^20^. Experiments were performed using P4-P7 passages, LEC identity was confirmed by Prox1 staining and only batches with over 95% Prox1-positive staining were used for experiments. For knockdown experiment, siRNA was used for knocking down *CXCR4* in Human dermal LECs in vitro. Cells were transfected with 50nM *siCXCR4* (ThermoFisher, catalog number 4390824) or negative control (ThermoFisher, catalog number AF4611) using Lipofectamine™ 2000 Transfection Reagent (Invitrogen, catalog number11668019) in Opti-MEM™ I Reduced Serum Medium (Gibco, Catalog number 31985062). After 6 hours of transfection, medium was changed with normal endothelial culture medium. The efficiency was evaluated after 48 hours transfection by using qPCR and western blot to check the expression level of *CXCR4* before subsequent experiments. For OSS, the human dermal LECs were exposed to OSS conditions (0.3 dynes/cm2) on a test tube rocker (Thermolyne Speci-Mix Aliquot Mixer Model M71015, Barnstead International) in a 37 °C incubator with 5% CO2. AKT activator SC-97 was administered 10 µM in HDLEC for 30 min.

### Immunofluorescence Staining and Confocal Microscopy

For whole-mount immunostaining, tissues were fixed in 2% paraformaldehyde (PFA) overnight at 4°C. Fixed tissues were washed with phosphate-buffered saline (PBS) and blocked in blocking buffer overnight at 4°C. Samples were then incubated with primary antibodies overnight at 4°C, followed by washing with PBST (PBS containing 0.1% Triton X-100). Appropriate secondary antibodies were applied for 2 hours at room temperature, after which tissues were washed with PBST and mounted using ProLong mounting medium (Sigma-Aldrich, P36931).

For cell immunostaining, cells were cultured on cover glasses in 24-well plates pre-coated with 0.1% gelatin overnight. Cells were transfected with siRNA at approximately 50% confluence. After 48 hours, cells were fixed with 2% PFA at room temperature and washed four times with PBS. Samples were blocked for 1 hour in blocking buffer, followed by incubation with primary antibodies for at least 2 hours at room temperature. Cells were subsequently washed with PBST and incubated with fluorophore-conjugated secondary antibodies diluted in blocking solution for 2 hours. After additional washes with PBST, samples were mounted using ProLong mounting medium with DAPI (Cell Signaling Technology, 8961S).

Microscopy images were acquired using a Nikon AX laser scanning confocal microscope mounted on an Eclipse Ti2 inverted microscope stand (Nikon Instruments). The system was equipped with a DUX-ST detector unit and LUD-S4 laser light source, and images were collected using NIS-Elements software (Nikon).

Detailed information on all primary and secondary antibodies used in this study is provided in Supplementary Table S1.

### Western Blot

To examine signaling changes in response to various treatments, cells were lysed in RIPA buffer (Invitrogen, PI89900) supplemented with a protease inhibitor cocktail (Pierce, A32959). Protein concentrations were determined using the BCA Protein Assay Kit (Thermo Fisher Scientific, 23227). Equal amounts of protein were subjected to SDS–PAGE, transferred to PVDF membranes, and analyzed by Western blotting. Protein bands were detected using SuperSignal™ West Pico Plus chemiluminescent substrate (Thermo Fisher Scientific) and imaged with the iBright™ 1500 Imaging System (Thermo Fisher Scientific). Band intensities were quantified using ImageJ (version 2.14.0). All raw data used for quantification are provided in the Supplementary Information, and details of primary and secondary antibodies are listed in Table S1.

### Quantitative Real-Time PCR (qRT–PCR)

Total RNA was extracted from human dermal LECs using TRIzol reagent (Thermo Fisher Scientific, 15596018) according to the manufacturer’s instructions. RNA concentration and purity were determined prior to downstream analysis.

Complementary DNA (cDNA) was synthesized from total RNA using the iScript cDNA Synthesis Kit (Bio-Rad, 1708891). Quantitative real-time PCR (qRT–PCR) was performed using Power SYBR Green PCR Master Mix (Applied Biosystems/Life Technologies) on a StepOnePlus Real-Time PCR System (Applied Biosystems). Each sample was analyzed individually in each qPCR run, and reactions were performed in technical replicates. Gene expression levels were normalized to the housekeeping gene GAPDH, and relative expression was calculated using the 2^−ΔΔCt^ method. Primer sequences used in this study are listed in Supplementary Table S2.

### Statistics

All data are presented as mean ± standard error of the mean (SEM) unless otherwise indicated. Statistical analyses were performed using GraphPad Prism. Comparisons between two groups were conducted using an unpaired two-tailed Student’s t-test, whereas comparisons among multiple groups were analyzed using one-way analysis of variance (ANOVA) followed by appropriate post hoc tests. The number of biological replicates (n) is indicated in the corresponding figure legends. Differences were considered statistically significant when P < 0.05. All experiments were repeated independently at least three times unless otherwise stated.

### Study Approval

All animal experiments were performed in accordance with the guidelines of the Institutional Animal Care and Use Committee (IACUC) of Temple University. All experimental protocols were reviewed and approved by the Temple University IACUC, and all procedures were conducted in compliance with the Guide for the Care and Use of Laboratory Animals published by the National Institutes of Health (NIH).

## Results

### CXCR4 is required for mesenteric lymphatic valve formation

To investigate the functional role of *CXCR4* in embryonic lymphatic valve development, we generated lymphatic endothelial cell (LEC)–specific *Cxcr4* conditional knockout mice by crossing *Cxcr4^f/f^* mice with *Flt4CreER^T2^* mice. Conditional deletion of *Cxcr4* in LECs was induced in pregnant females by administering tamoxifen (TM; 5 mg/40 g body weight) at embryonic days E13.5 and E14.5, coinciding with the onset of lymphatic valve formation as shown in Fig. 1A. Embryonic mesenteric lymphatic vessels were collected at E18.5 for analysis. Mesenteric lymphatic vessels from *Flt4CreER^T2^, Cxcr4^f/f^* embryos displayed a marked reduction in lymphatic valves, as indicated by decreased numbers of Prox1^high^ and FOXC2^high^ valve endothelial cells compared with controls (Fig. 1B-C). Furthermore, VEGFR3 staining also showed the defects of lymphatic valve in *Flt4CreER^T2^, Cxcr4^f/f^* embryos (Fig. 1D). Together, these findings demonstrate that CXCR4 signaling is required for proper lymphatic valve formation during embryonic lymphatic vascular development.

**Figure 1.**
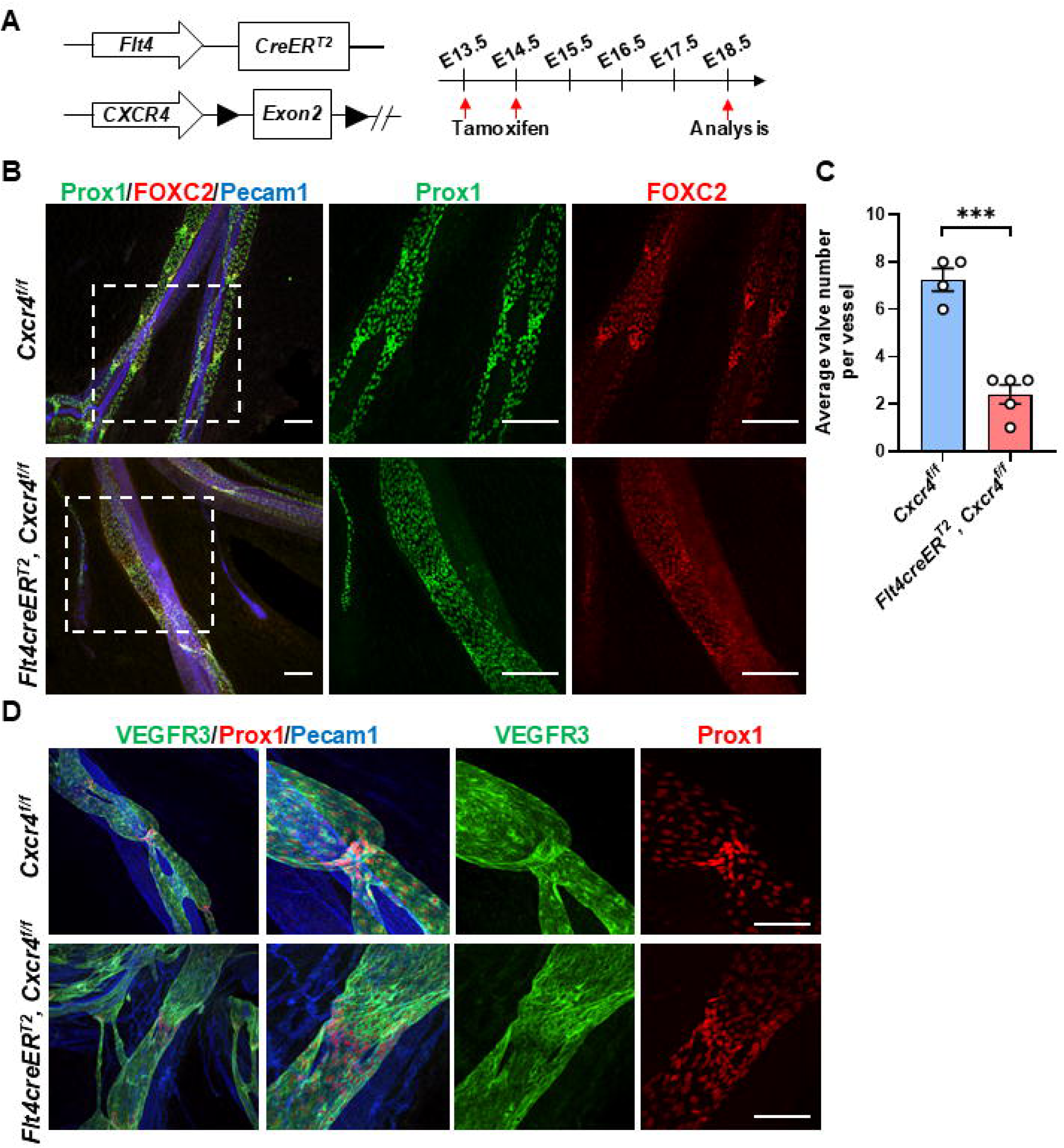
Lymphatic endothelial cell-specific deletion of Cxcr4 impairs embryonic lymphatic valve formation. **A.** Diagram of mouse model and tamoxifen administration. **B.** Whole mount staining for Prox1, FOXC2, and Pecam1 in mesentery of E18.5 embryos. **C.** The quantification of lymphatic valves (numbers per vessle) between *Cxcr4^f/f^* and *Flt4CreER^T2^, Cxcr4^f/f^* embryos, N=4-5. **D.** Whole mount staining for Prox1, VEGFR3, and Pecam1 in mesentery of E18.5 embryos. Scale bar: 100 µm in B and 50 µm in D. Data are mean± s.e.m. \*\*\**P*<0.0001, unpaired two-tailed Student’s *t*-test.

### CXCL12 is required for lymphatic valve formation

To determine whether CXCL12 similarly contributes to lymphatic valve development, we next examined lymphatic vessels in *Cxcl12−/−* embryos. Immunofluorescence analysis of embryonic mesenteric lymphatic vessels revealed a marked reduction in lymphatic valve structures in *Cxcl12−/−* embryos compared with wild-type and heterozygous littermates (Fig. 2A). Specifically, the number of PROX1^high^ valve LECs clusters was substantially decreased in the mesenteric collecting vessels of *Cxcl12−/−*embryos. Quantitative analysis confirmed a significant reduction in valve density, measured as the number of PROX1^high^ valves per millimeter of lymphatic vessel length (Fig. 2B). Despite this reduction in valve formation, the overall morphology and patterning of the mesenteric lymphatic vessel network appeared largely preserved. These findings indicate that CXCL12 is required for proper lymphatic valve development, supporting a model in which CXCL12/CXCR4 signaling regulates the formation of lymphatic valves during embryonic lymphatic vascular development.

**Figure 2.**
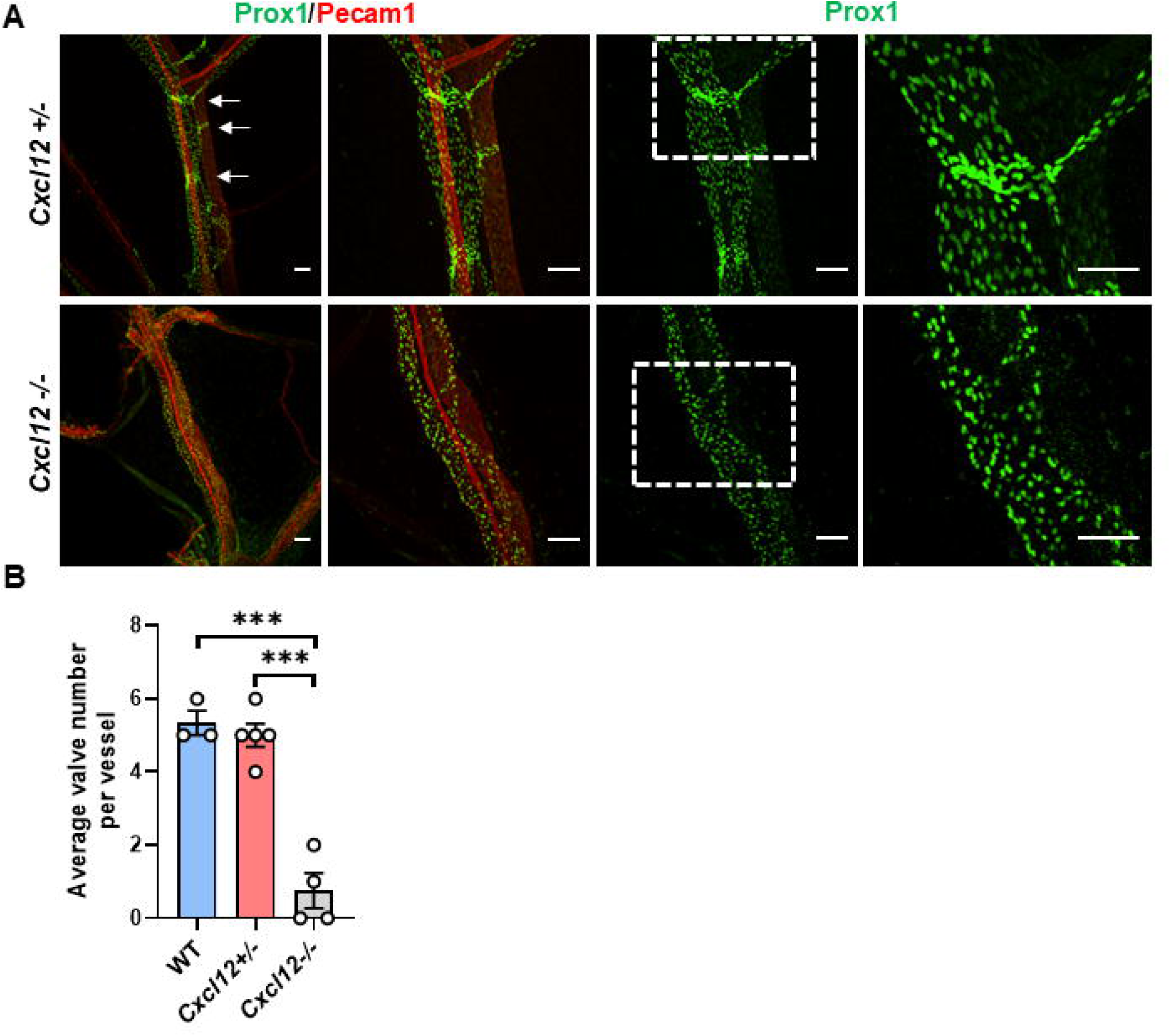
*Cxcl12* deficiency impairs embryonic lymphatic valve numbers. **A.** Whole mount staining for Prox1 and Pecam1 in mesentery of E17.5 embryos. White arrows indicate mesentery lymphatic valve. **B.** The quantification of lymphatic valves between wild type (WT), *Cxcl12+/-*, and *Cxcl12−/−* embryos, N=3-5. Scale bar: 100 µm. Data are mean± s.e.m. \*\*\*\**P*<0.0001, one-way ANOVA followed by Turkey’s test.

### Blood endothelial cell derived CXCL12 is required for lymphatic valve formation

Our previous work demonstrated that CXCR4 ligand CXCL12 derived from neurons plays a critical role in the embryonic dermal lymphatic development ^13^. However, the cellular sources of CXCL12 in the mesentery, where lymphatic valves form extensively during embryonic development remain unknown. To identify the source of CXCL12, we utilized *Cxcl12-DsRed* KIKO reporter mice, in which CXCL12 expression is reported by RFP. Whole mount staining revealed that RFP expression colocalized with the blood vessel marker connexin 40 (CX40), indicating that CXCL12 is predominantly expressed by mesenteric blood vessels (Fig. 3A). In contrast to the dorsal skin, where CXCL12 is primarily produced by neurons, RFP expression did not colocalize with the neuronal marker Tuj1 in the mesentery, further supporting the conclusion that blood vessels are major source of mesenteric CXCL12 (Fig. 3B). To determine whether blood endothelial cell derived CXCL12 contributes to lymphatic valve formation, we generated endothelial cell (EC)–specific *Cxcl12* conditional knockout mice by crossing *Cxcl12^f/f^* mice with *Cdh5CreER^T2^* mice. Using the same strategy described in Fig. 1A, we analyzed lymphatic valve numbers and morphology in the embryonic mesentery. Compared with littermate controls, *Cdh5CreER^T2^, Cxcl12^f/f^* embryos exhibited decreased lymphatic valve number, accompanied by a reduction in Prox1^high^ and FOXC2^high^ valve endothelial cells (Fig. 3C-E). Collectively, these results indicate that mesenteric blood vessel derived CXCL12 is required for normal lymphatic valve formation, suggesting a potential paracrine signaling from blood vessels to lymphatic vessels in the mesentery that may coordinate mesenteric lymphatic valve development.

**Figure 3.**
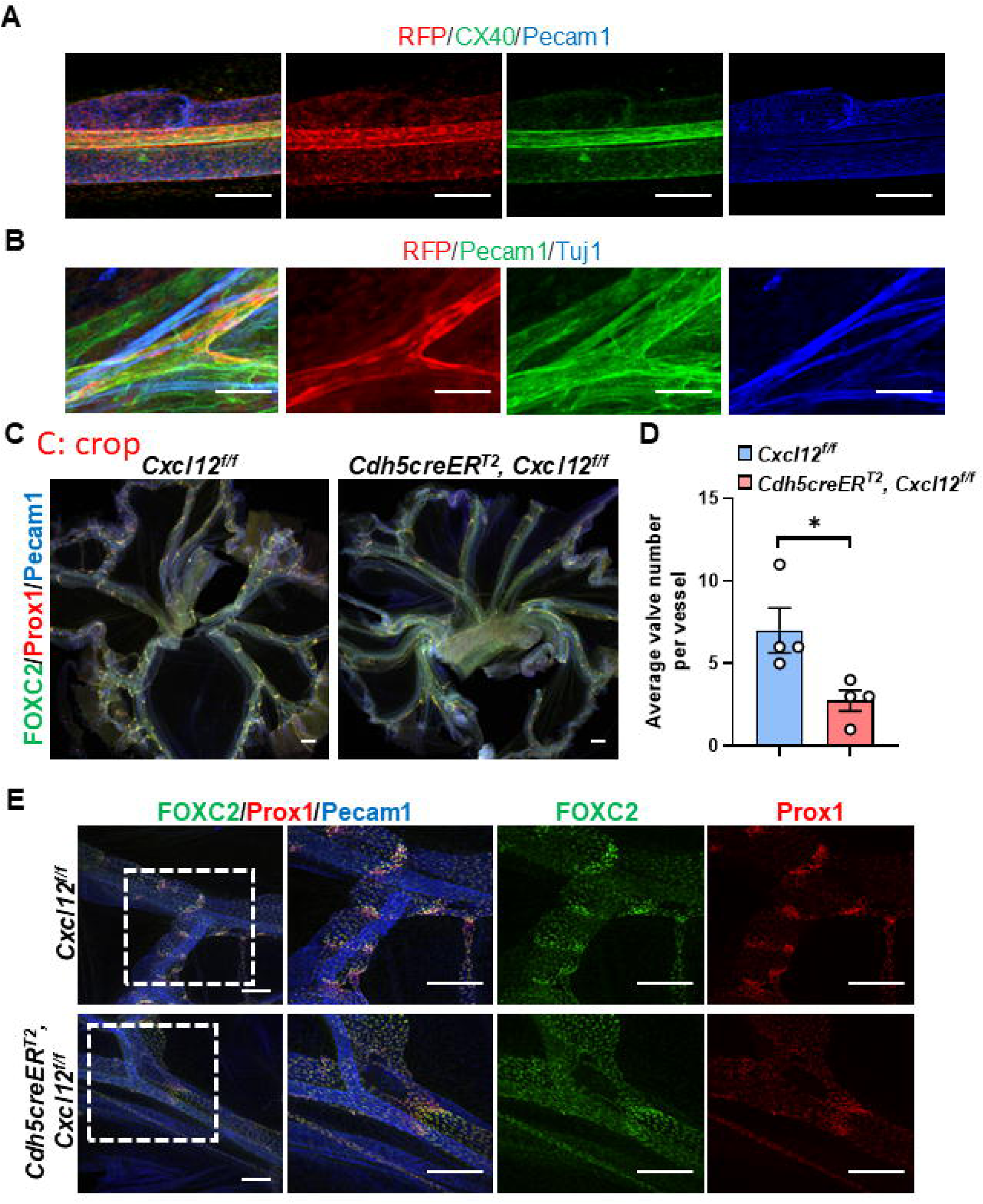
Endothelial cell-derived CXCL12 is required for embryonic lymphatic valve formation. **A.** Whole mount staining for CX40, RFP, and Pecam1 in mesentery of E14.5 embryos. **B.** Whole mount staining for Tuj1, RFP, and Pecam1 in mesentery of P5 pups. **C.** Whole mount staining for Prox1, FOXC2, and Pecam1 in mesentery of E18.5 embryos. **D.** The quantification of lymphatic valves between *Cxcl12^f/f^* and *Cdh5CreER^T2^, Cxcl12^f/f^* embryos, N=4. **E.** Whole mount staining for Prox1, FOXC2, and Pecam1 in mesentery of E18.5 embryos. Scale bar: 100 µm in A, B and E, 200 µm in C. Data are mean± s.e.m. \**P*<0.05, unpaired two-tailed Student’s *t*-test.

### CXCR4 Knockdown Inhibits OSS-Induced valve gene expression through AKT signaling

To further elucidate the underlying molecular mechanisms by which CXCL12/CXCR4 regulates lymphatic valve formation, we performed RNA sequencing (RNA-seq) analysis in human dermal LECs treated with stromal cell-derived factor 1 (SDF-1, also known as CXCL12) or subject to siRNA-mediated knockdown of *CXCR4*. Differential gene expression analysis was conducted to identify genes that were upregulated upon SDF1 stimulation and downregulated following *CXCR4* knockdown, and the overlapping gene set was subsequently analyzed. Gene ontology and pathway analyses revealed enrichment of pathways associated with FOXO1 signaling, lymphatic valve formation, and fluid shear stress in SDF1 treatment group. In contrast, *CXCR4* knockdown downregulated pathways related with lymphatic valve formation, TGFβ signaling pathway, and sprouting angiogenesis (Fig. 4A-B). Comparing these datasets identified a total of 64 overlapping genes that were positively regulated by CXCL12 and negatively regulated by *CXCR4* knockdown (Fig. 4C). Notably, this gene set included several established key regulators of lymphatic valve formation and maturation, including *FOXC2* and *GJA4*, suggesting that they function downstream of CXCL12/CXCR4 signaling. This finding suggests that CXCL12/CXCR4 potentially contributes to the control of lymphatic valve formation. Because oscillatory shear stress is a key biomechanical stimulus that drives lymphatic valve development through induction of PROX1, FOXC2, and GATA2 ^6–8, 21^, we next examined whether CXCR4 participated in flow dependent valve signaling. Exposure of LECs to OSS significantly increased FOXC2 protein expression, confirming activation of the valve formation program (Fig. 4D-E) and is consistent with previous reports that FOXC2 is induced by OSS. Interestingly, OSS also increased CXCR4 protein levels, which indicates that CXCR4 might be regulated by lymph flow.

**Figure 4.**
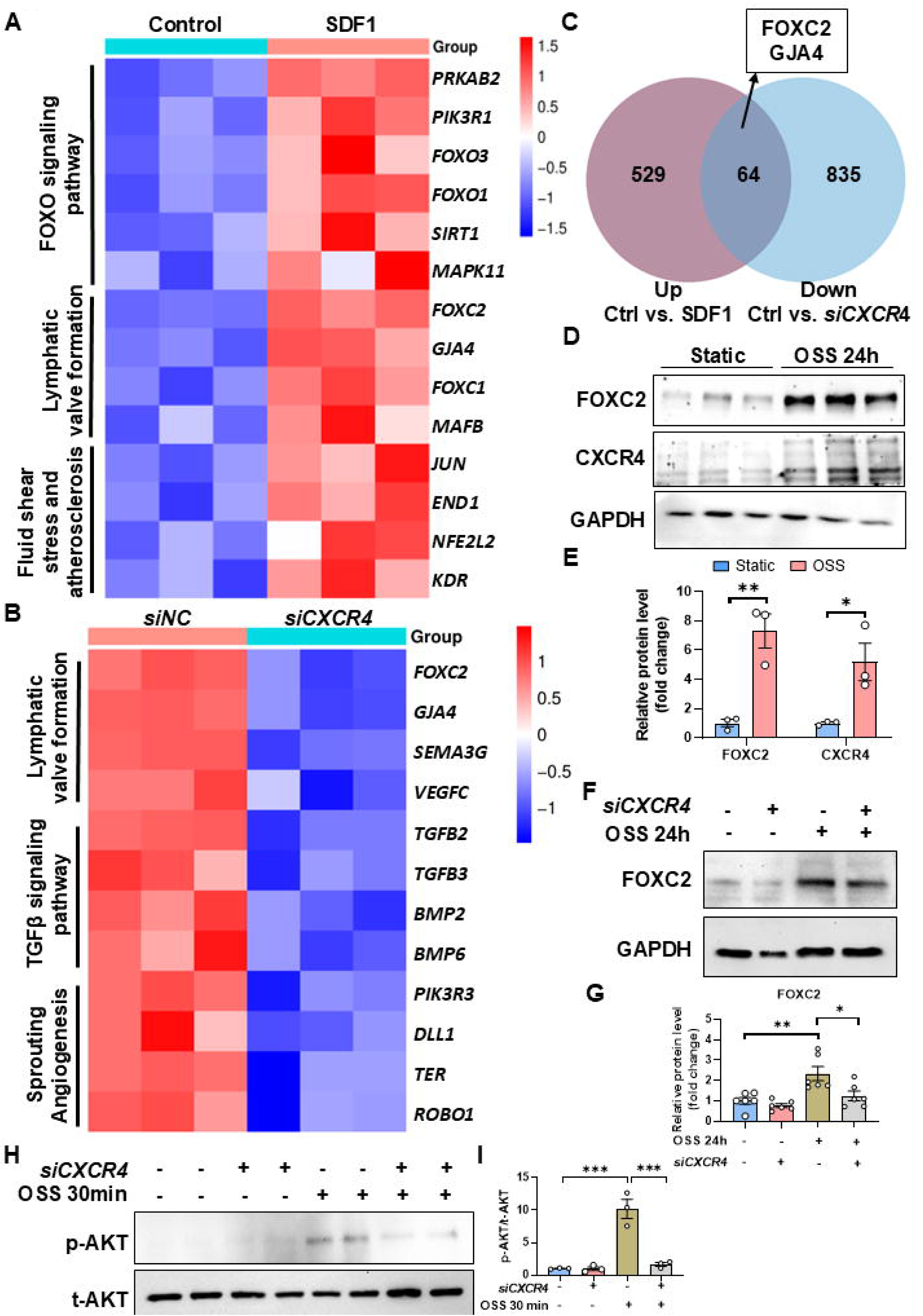
CXCR4 deficiency impairs OSS-induced upregulation of valve formation related genes via AKT signaling pathway. **A.** Heatmap of upregulated pathways and the involved genes in SDF1 treatment group compared with control in Human dermal LECs. **B.** Heatmap of downregulated pathways and the involved genes in *siCXCR4* treatment group compared with control in Human dermal LECs. **C.** Overlapped genes of upregulated in SDF1 treatment and downregulated in *siCXCR4* treatment. **D.** Western blot targeted FOXC2 and CXCR4 under static and OSS condition for 24h. **E.** The quantification of FOXC2 and CXCR4 intensities in D. **F.** Western blot targeted FOXC2 treated with *siCXCR4* and OSS for 24h. **G.** The quantity of FOXC2 intensity. **H.** Western blot targeted p-AKT and t-AKT treated with *siCXCR4* and OSS for 30min. **G.** The quantity for the ratio of p-AKT/ t-AKT intensity. Data are mean± s.e.m. \**P*<0.05, \*\**P*<0.01, \*\*\**P*<0.001, unpaired two-tailed Student’s *t*-test (E) and one-way ANOVA followed by Turkey’s test (G, I). n number and experimental repeats

Given the defective lymphatic valve formation and reduced PROX1 and FOXC2 expression observed in *Flt4CreER^T2^, Cxcr4^f/f^* embryonic mesenteries (Fig. 1B), we next investigated whether CXCR4 is required for OSS induced valve gene expression in vitro. Human dermal LECs were transfected with *siCXCR4* or scrambled control siRNA and subsequently exposed to static or OSS conditions. Under OSS, *CXCR4* knockdown markedly attenuated induction of FOXC2 protein levels (Fig. 4F-G). Consistently, q-RTPCR analysis demonstrated significantly reduced expression of *FOXC2* and *GJA4* in siCXCR4 treated LECs exposed to OSS (S-Fig1. A-B).

Because AKT signaling has been reported to be a key mediator of OSS-induced valve signaling ^15,16^, we next assessed AKT phosphorylation (p-AKT). OSS robustly increased p-AKT levels in control cells, whereas *siCXCR4 treated LECs* abolished this response (Fig. 4H-I). These results indicate that CXCR4 functions upstream of AKT in the mechanotransduction pathway that links oscillatory lymph flow to valve gene activation, revealing a critical role for CXCR4 in translating mechanical cues into transcriptional programs necessary for lymphatic valve formation.

### Embryonic deletion of *Cxcr4* in lymphatic endothelial cells promotes FOXO1 nuclear retention

FOXO1 is a critical downstream effector of AKT signaling and a key regulator of lymphatic valve formation ^17^. In LECs, activation of the PI3K/AKT pathway promotes phosphorylation of FOXO1, leading to its cytoplasmic retention and inhibition of its transcriptional activity. In contrast, reduced AKT signaling leads to the accumulation of unphosphorylated FOXO1 in the nucleus, where it represses the expression of genes required for lymphatic valve development ^17^. Previous studies have shown that excessive nuclear FOXO1 activity suppresses the expression of critical valve regulators, including PROX1, FOXC2, and GATA2, thereby impairing lymphatic valve specification and maturation. Given that *CXCR4* knockdown attenuated AKT activation in response to oscillatory shear stress, we next examined whether CXCR4 regulates FOXO1 activity. Consistent with reduced AKT signaling, knockdown of *CXCR4* in LECs decreased FOXO1 phosphorylation and enhanced nuclear FOXO1 localization (Fig. 5A-D). To determine whether this mechanism also occurs in vivo, we analyzed nuclear FOXO1 expression in mesenteric lymphatic vessels in *Flt4CreER^T2^, Cxcr4^f/f^* embryos. Compared with littermate controls, *Flt4CreER^T2^, Cxcr4^f/f^* embryos exhibited significantly increased nuclear FOXO1 accumulation within lymphatic valve regions (Fig. 5E-F). Together, these findings indicate that CXCL12/CXCR4 signaling promotes AKT-dependent phosphorylation and inhibition of FOXO1, thereby preventing its nuclear accumulation during lymphatic valve development.

**Figure 5.**
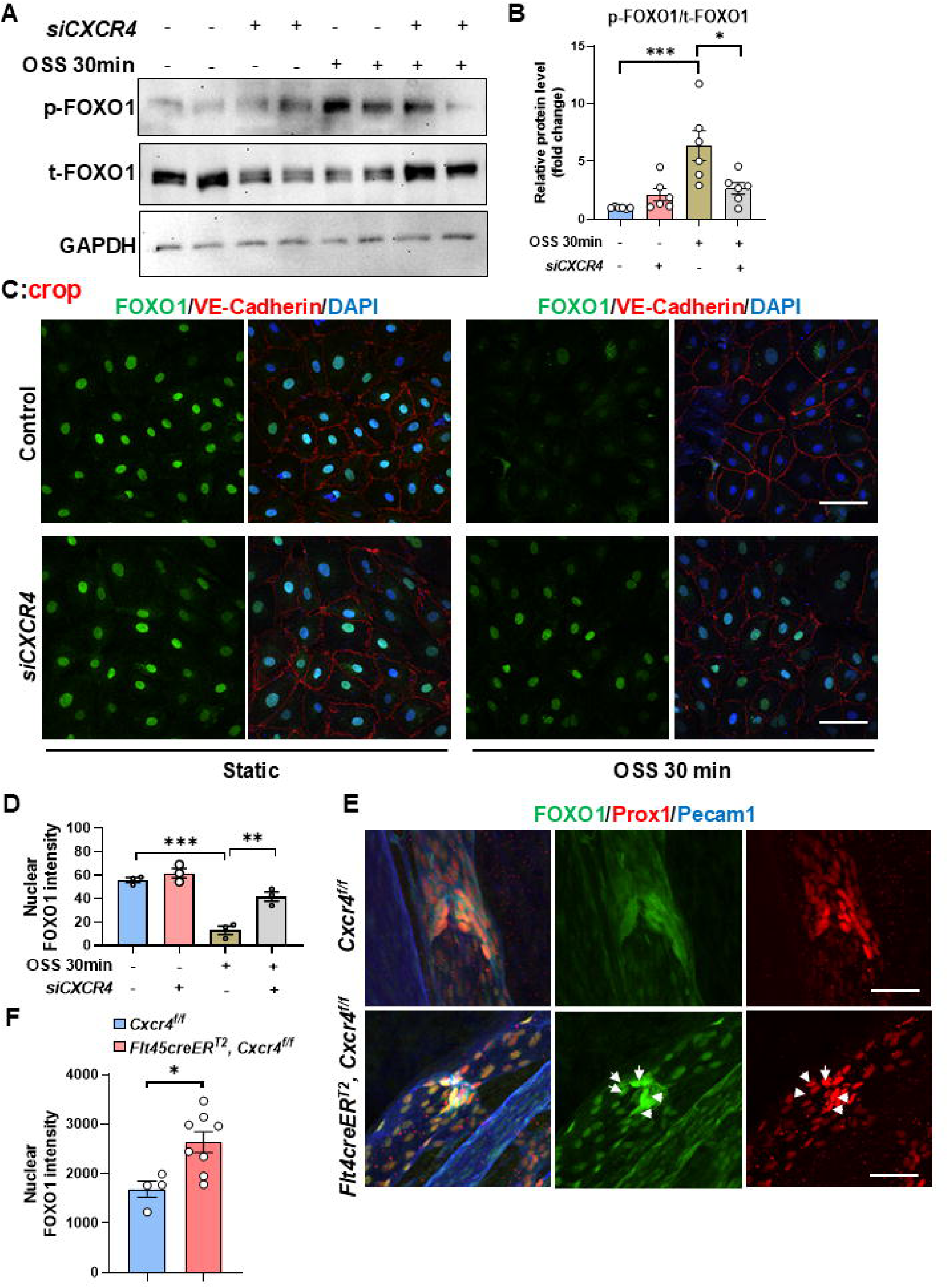
CXCR4 deficiency results in nuclear FOXO1 retention in lymphatic endothelial cells. **A.** Western blot targeted p-FOXO1 and t-FOXO1 under static and OSS condition for 30min. **B.** The quantity for the ratio of p-FOXO1/ t-FOXO1 intensity. **C.** Immunofluorescence for FOXO1, VE-Cadherin and DAPI in HELECs with the treatment of siCXCR4 and OSS for 30 min. **D.** The quantification of FOXO1 intensity in the nuclear in Human dermal LECs, N=3. **E.** Whole mount staining for FOXO1, Prox1, and Pecam1 in mesentery of E18.5 embryos. **F.** The quantification of FOXO1 between *Cxcr4^f/f^* and *Flt4CreER^T2^, Cxcr4^f/f^* embryos, N=4-8. Scale bar: 100 µm. Data are mean± s.e.m. \**P*<0.05, \*\*\**P*<0.001, unpaired two-tailed Student’s *t*-test (D, F) and one-way ANOVA followed by Turkey’s test (B).

### AKT activation rescues lymphatic valve formation in *Flt4CreER^T2^, Cxcr4^f/f^* embryos

Given that AKT signaling is a key regulator of lymphatic valve formation, we next tested whether pharmacological activation of AKT could rescue the valve defects observed in *Flt4CreER^T2^, Cxcr4^f/f^* embryos. Pregnant females were administered the AKT activator SC-79 at E15.5 after tamoxifen administration. Notably, SC-79 treatment significantly restored lymphatic valve formation in the mesenteric collecting vessels of *Flt4CreER^T2^, Cxcr4^f/f^* embryos (Fig. 6A-B). To determine the underlying mechanism, we examined FOXO1 activity following AKT activation. SC-79 administration increased FOXO1 phosphorylation, which promotes its cytoplasmic retention and suppresses its transcriptional repressor activity (Fig. 6C-E). Consistent with this mechanism, nuclear localization of FOXO1 was significantly reduced in human dermal LECs following SC-79 treatment (Fig. 6F-G). These findings indicate that AKT activation rescues lymphatic valve defects in *Flt4CreER^T2^, Cxcr4^f/f^* embryos by regulating FOXO1 phosphorylation and subcellular localization, supporting a model in which CXCR4 promotes lymphatic valve development through the AKT/FOXO1 signaling axis (Fig. 7).

**Figure 6.**
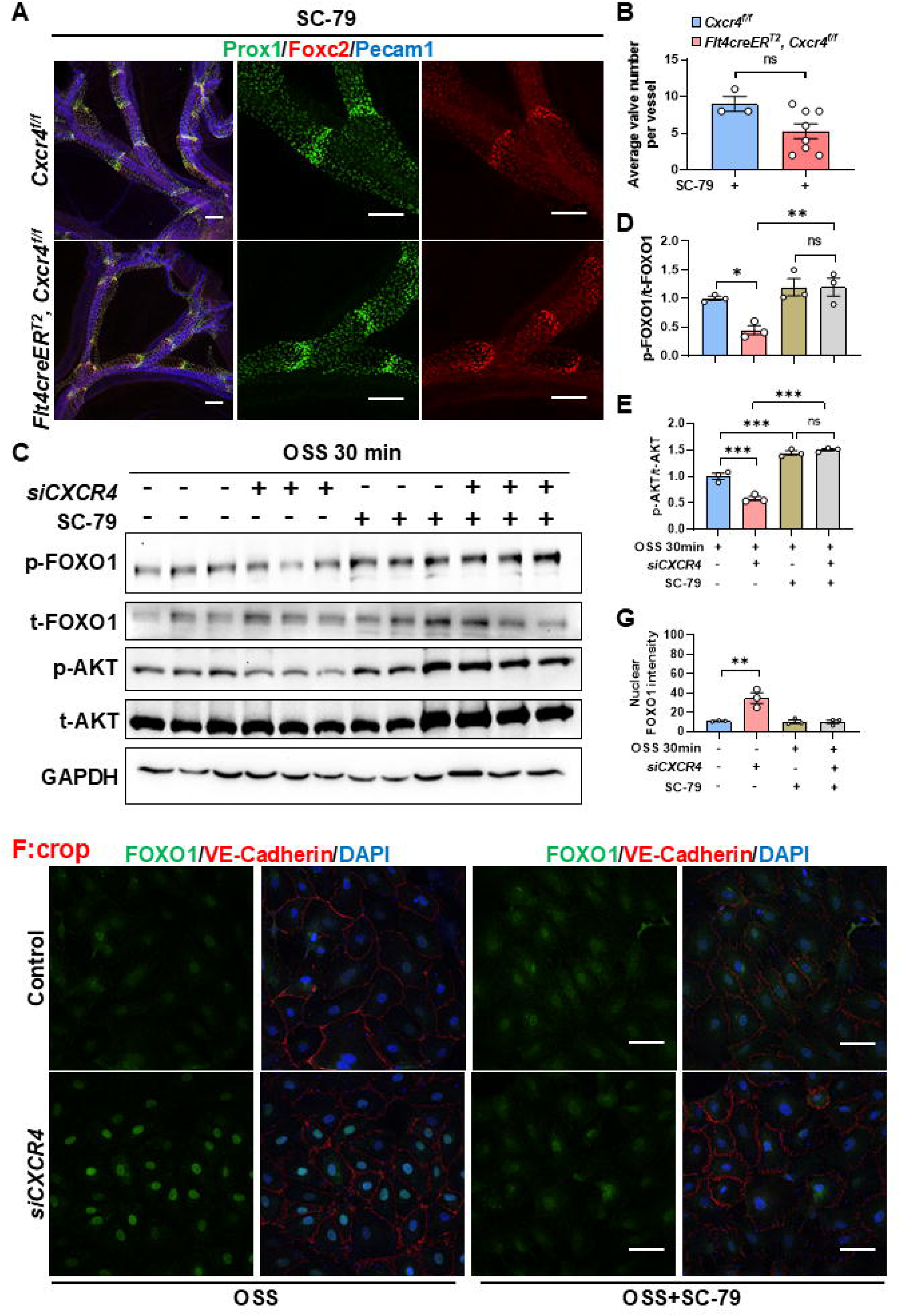
AKT activation rescues lymphatic valve defects in *Flt4CreER^T^*^2^*, Cxcr4^f/f^* embryonic mice. **A.** Whole mount staining for FOXC2, Prox1, and Pecam1 in mesentery of E18.5 embryos treated with tamoxifen at E13.5 and half does at E14.5, and AKT activator SC-79 at E15.5. **B.** The quantification of lymphatic valves between *Cxcr4^f/f^* and *Flt4CreER^T2^, Cxcr4^f/f^* embryos, N=3-8. **C.** Western blot targeted p-FOXO1, t-FOXO1, p-AKT and t-AKT treated with *siCXCR4* and OSS for 30 min. **D-E.** The quantity for the ratio of p-FOXO1/ t-FOXO1 and p-AKT/ t-AKT intensity. **F.** Immunofluorescence for FOXO1, VE-Cadherin and DAPI in human dermal LECs with the treatment of pretreating with SC-79 for 6h, *siCXCR4*, and OSS 30 min. **G.** The quantification of FOXO1 in nuclear, N=3. Scale bar: 100 µm. Data are mean± s.e.m. \**P*<0.05, \*\**P*<0.01, \*\*\**P*<0.001, unpaired two-tailed Student’s *t*-test (B, G) and one-way ANOVA followed by Turkey’s test (D, E).

**Figure 7.**
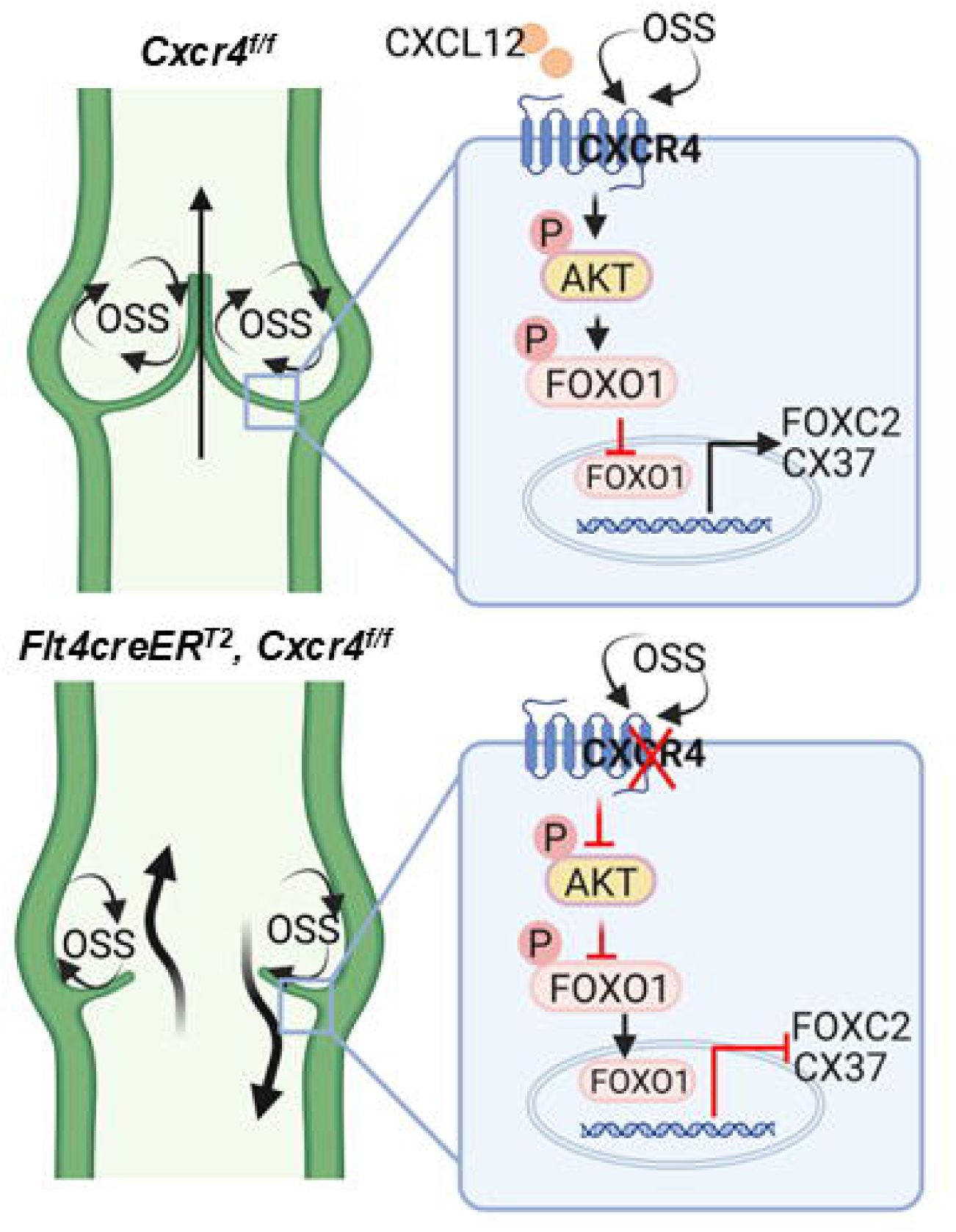
CXCR4 controls lymphatic valve formation via AKT-FOXO1 signaling pathway. Schematic illustration shows that CXCR4 deficiency compromises lymphatic valve formation, in which the response to OSS induced FOXC2 expression is damaged. Mechanically, CXCR4 knockdown inhibits AKT phosphorylation and its downstream p-FOXO1, which makes more FOXO1 retention in nuclear and represses valve formation genes like FOXC2 and CX37 expression.

## Discussion

Efficient lymphatic function relies on the precise formation and maintenance of lymphatic valves; despite these key functions, the molecular pathways integrating biochemical cues with mechanotransduction in LECs remain incompletely understood. Here, we identify the CXCL12/CXCR4 axis as a central regulator of lymphatic valve development, acting through AKT-mediated inhibition of the transcriptional repressor FOXO1. Using LEC-specific *Cxcr4* knockout and *Cxcl12*-deficient embryos, we demonstrate that loss of CXCL12/CXCR4 signaling leads to a profound reduction in embryonic mesenteric lymphatic valves. Mechanistically, CXCR4 is required for AKT activation in response to oscillatory shear stress. OSS phosphorylates FOXO1 to promote its cytoplasmic retention, thereby relieving repression of valve-specific transcriptional programs, including PROX1 and FOXC2. Importantly, pharmacological activation of AKT rescues FOXO1 phosphorylation and restores valve formation in *Flt4CreER^T2^, Cxcr4^f/f^* embryos, confirming that AKT/FOXO1 signaling acts downstream of CXCR4 in lymphatic mechanotransduction. Our findings reveal a previously unrecognized mechanochemical crosstalk, whereby LECs chemokine signaling converges with flow-dependent pathways to orchestrate valve endothelial cell specification and morphogenesis. This work expands the functional repertoire of CXCL12/CXCR4 beyond lymphatic migration and patterning, highlighting its role in coordinating the transcriptional and mechanical inputs that govern valve formation. Clinically, these insights provide a mechanistic basis for congenital lymphatic valve defects and lymphedema associated with impaired chemokine signaling.

Recent studies revealed mechanical sensors such as Piezo1 and YAP/TAZ play vital roles in lymphatic valve formation ^22–26^. For example, in mouse embryos, deletion of *Yap/taz* in LECs has no effects on the initiation of lymphatic vessel formation but causes valve degradation over time as they are required to maintain Prox1 expression. Our results (Fig. 4D) showed that OSS increased CXCR4 protein level, which means mechanical forces from lymph flow influence CXCR4 level. In future studies, it will be necessary to determine how CXCL12/CXCR4 interacts with other mechanosensitive pathways like YAP/TAZ or Piezo1, and whether modulation of this axis could be harnessed to therapeutically enhance lymphatic valve regeneration.

We previously reported that Schwann cell-derived CXCL12 plays a critical role in the initiation of embryonic lymphatic development ^13^. However, the cellular sources of CXCL12 in mesenteric lymphatics have remained largely undefined. In the present study, we demonstrate that CXCL12 is abundantly expressed in the embryonic mesentery, with particularly high levels coming from blood vessels. The crosstalk between lymphatic vessels and blood vessels suggest that blood endothelial–derived CXCL12 is key to support valve formation in mesentery lymphatics. These findings raise the possibility that paracrine sources of CXCL12, in different organs might play an important role in organ-specific lymphatic valve formation. Future studies will be required to define the relative contributions of these cellular sources and to elucidate the crosstalk between blood vessels, stromal cells, and lymphatic vessels that coordinate lymphatic vascular morphogenesis.

In conclusion, our study establishes CXCL12/CXCR4/AKT/FOXO1 signaling as a critical mechano-chemical axis driving lymphatic valve formation, integrating extracellular cues with transcriptional regulation to ensure proper lymphatic vascular development.

## Author Contributions

Conceptualization, L.D, J.P. and X.L.; study design, L.D., J.P. and X.L.; performed experiments, L.D. and J.P.; mouse experiments and tissue analysis, L.D.; imaging and quantitative analysis, L.D. and J.P.; molecular experiments, J.P. and L.D.; data analysis and interpretation, L.D., J.P. and X.L.; writing—original draft preparation, J.P.; writing—review and editing, J.P., L.D., E.D, L.F., H.L. and X.L.; supervision, X.L.; funding acquisition, X.L.. All authors have read and agreed to the published version of the manuscript.

## Acknowledgements

We thank Dr. Ralf Admas for the source of *Cdh5creER^T2^* mice. We thank Dr. Sagrario Ortega for the source of *Flt4creER^T2^* mice. We thank Lemole Center Imaging Core for providing advanced microscope and equipment support in this study. Graphic image was created by using Biorender.com.

## Non-standard Abbreviations and Acronyms

CXCL12: Chemokine (C-X-C Motif) Ligand 12
CXCR4: C-X-C Chemokine Receptor Type 4
CX4: Connexin 40
E: Embryonic day
FOXO1: Forkhead Box Protein O1
FOXC2: Forkhead Box C2
GATA2: GATA-binding protein 2
LECs: Lymphatic endothelial cells
OSS: Oscillatory shear stress
TM: Tamoxifen
VEGFR3: Vascular Endothelial Growth Factor Receptor 3

