## supplementary file for "CXCL12/CXCR4 Governs Lymphatic Valve Formation Through Flow Dependent AKT/FOXO1/FOXC2 Activation"

*The authors contribute equally to this manuscript

To whom all correspondence should be addressed:

Xiaolei Liu, Ph.D.

Temple University School of Medicine

Room 1045B MERB

3500 North Broad Street

Philadelphia, PA, 19140

Short title: CXCR4 potentiates flow dependent valve formation

Total word: 5539

Table S1.

| **Target antigen** | **Vendor** | **Catalog Number** | **Working Concentration** |
| --- | --- | --- | --- |
| VEGFR3 | R and D systems | AF1002 | IF 1:500 |
| FOXC2 | R and D systems | AF6989 | IF 1:500 |
| FOXO1 | CST | 2880S | WB 1:1000  IF 1:500 |
| Phospho-FOXO1 | CST | 2486S | WB 1:1000 |
| Prox1 | Abcam | ab101851 | IF 1:500 |
| Pecam1 | BD Bio | 557355 | IF 1:500 |
| VE-Cadherin | R and D systems | AF1002 | IF 1:1000 |
| Phospho-Akt | CST | 9271S | WB 1:1000 |
| Total-Akt | CST | 9272S | WB 1:1000 |
| RFP | Rockland | 600-406-379 | IF 1:1000 |
| Tuj1 | Biolegend | 801202 | IF 1:1000 |
| GAPDH | ThermoFisher | MA5-15738-HRP | WB 1:5000 |
| Donkey anti-rat, Alexa Fluor™ 488 | ThermoFisher | A-21208 | IF 1:1000 |
| Donkey anti-goat, Alexa Fluor™ 488 | ThermoFisher | A-11055 | IF 1:1000 |
| Donkey anti-goat, Alexa Fluor™ 594 | ThermoFisher | A-11058 | IF 1:1000 |
| Donkey anti-rabbit, Alexa Fluor™ 647 | Thermofisher | A-31573 | IF 1:1000 |
| Donkey anti-Rabbit Alexa Fluor™ 594 | Invitrogen | A-21207 | IF 1:1000 |
| Donkey anti-goat, Alexa Fluor™ 647 | Invitrogen | A-21447 | IF 1:1000 |
| CY5 donkey anti-Rabbit | Jackson Immuno | 711-175-152 | IF 1:1000 |
| CY3 donkey anti Rat | Jackson Immuno | 112-165-003 | IF 1:1000 |
| Cy5 donkey anti-Mouse | Jackson Immuno | 715-175-150 | IF 1:1000 |
| CY5 donkey anti-Rat | Jackson Immuno | 712-175-150 | IF 1:1000 |

Table S2

| **Gene** | **Species** | **Forward (5’-3’)** | **Reverse (5’-3’)** |
| --- | --- | --- | --- |
| *FOXC2* | Human | TCACCTTGAACGGCATCTACCAG | TGACGAAGCACTCGTTGAGCGA |
| *GJA4* | Human | TGCAAGAGTGTGCTAGAGGC | ACAAAGCAGTCCACGAGGTAG |
| *GAPDH* | Human | GGTGTGAACCATGAGAAGTATGA | GAGTCCTTCCACGATACCAAAG |

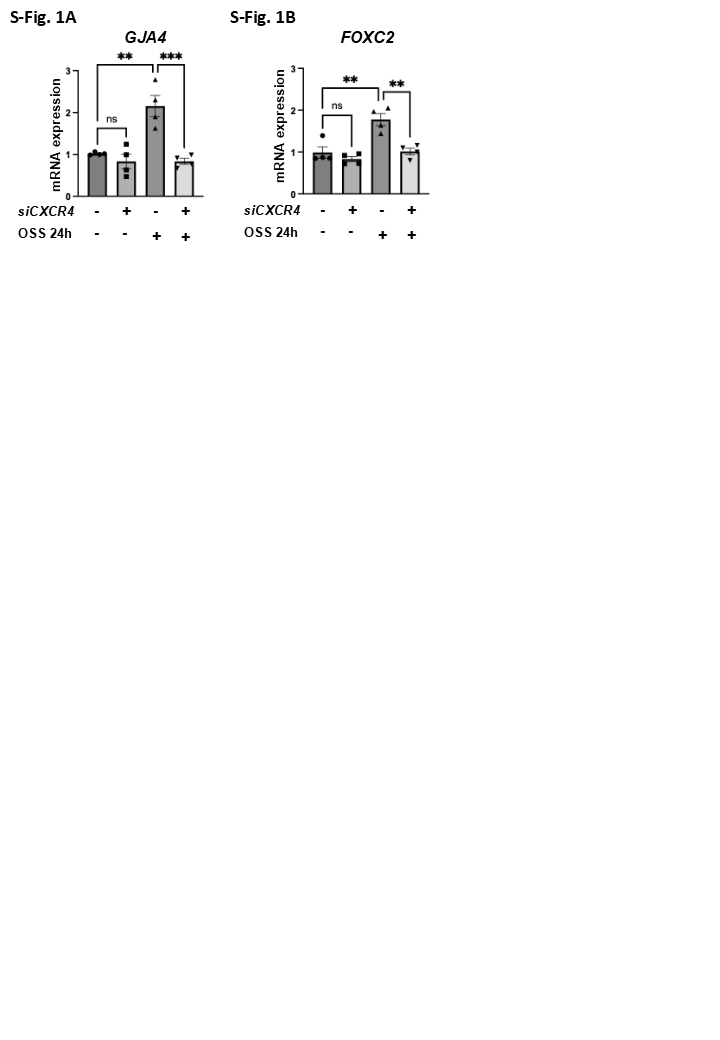

**Supplemental Figure 1. CXCR4 knockdown inhibits OSS-induced *GJA4* and *FOXC2* mRNA levels. A-B.** Human dermal LECs were transfected with *siCXCR4* or *siControl* and then exposed to OSS for 24h. qRT-PCR was used to check the mRNA levels of *GJA4* and *FOXC2*. Data are mean± s.e.m. ***P*<0.01, ****P*<0.001, one-way ANOVA followed by Turkey’s test (A, B).
